# Hidden in the deep: distinct benthic trajectories call for monitoring of mesophotic reefs

**DOI:** 10.1101/2021.08.01.454664

**Authors:** A Hernandez-Agreda, FM Sahit, N Englebert, O Hoegh-Guldberg, P Bongaerts

## Abstract

Long-term monitoring studies are central to coral reef ecology and conservation management. However, ongoing monitoring programs are almost exclusively focused on shallow depths, and it remains unclear to what extent those are representative of the whole ecosystem. Here, we present a temporal comparison (2012-2017) of directly adjacent shallow and mesophotic benthic communities across seven sites from the Great Barrier Reef and Western Coral Sea. We found a positive correlation initially between shallow and mesophotic coral cover, with higher cover at shallow depths. However, this correlation broke down after multiple disturbances, with coral cover declining only at shallow depths. Point-based tracking revealed the dynamic nature of mesophotic communities, with their consistent coral cover reflecting a net balance between substantial growth and mortality. Overall, the divergent trajectories highlight the urgency to expand monitoring efforts into mesophotic depths, to decipher the processes governing these habitats and enable better-informed management of the overall ecosystem.

## Introduction

Tropical coral reef ecosystems have undergone an unprecedented and accelerated decline resulting from escalating local and climatic stressors (Hughes et al., 2017; Jackson et al., 2014). Long-term monitoring programs play an integral role in documenting these changes and identifying the underlying mechanisms of decline by discriminating between natural and anthropogenic variability in ecosystem dynamics (De’ath et al., 2012; Hughes & Connell, 1999). For example, the basin-wide collapse of Caribbean reefs was recorded through such monitoring efforts, attributing the decline to the cumulative stress of acute (e.g. disease outbreaks, hurricanes, and thermal bleaching) and chronic disturbances (e.g. overfishing, tourism, and pollution) (Gardner et al., 2003; Jackson et al., 2014). While long-term monitoring programs are fewer in the Indo-Pacific, comprehensive monitoring on the Great Barrier Reef has documented the detrimental impacts of cyclones, outbreaks of crown-of-thorns starfish, and thermal bleaching, highlighting the extensive variability in trajectories at both small and regional spatial scales (De’ath et al., 2012; Hughes et al., 2017; Osborne et al., 2011). Long-term ecological monitoring is vital to improve our understanding of spatio-temporal patterns and processes and subsequently forms the basis of sound conservation management.

Mesophotic coral ecosystems (MCEs) are a major component of the global tropical reef habitat (~50-80%, Harris et al., (2013); Pyle & Copus, (2019)), yet they remain largely unsurveyed and therefore ignored in conservation planning (Bridge et al., 2013; Eyal et al., 2021; Rocha et al., 2018). Given their distinct environmental and biological characteristics compared to shallow reefs, MCEs have the potential to have distinct demographic characteristics (Kramer et al., 2020; Vermeij & Bak, 2003) and be differentially affected by disturbances (Frade et al., 2018; Smith et al., 2019). However, as coral reef monitoring is almost exclusively conducted at shallow depths (with the majority <15 m depth, (De’ath et al., 2012; Gardner et al., 2003)), the past and current trajectories of MCEs remain largely unknown (Bongaerts & Smith, 2019). Repeated surveys in the Caribbean have documented the possibility of substantial declines in coral cover at upper mesophotic depths (30-40 m) due to storms, thermal bleaching, and lionfish invasion (Hughes & Tanner, 2000; Lesser & Slattery, 2011). They have also highlighted how responses to disturbances and overall trajectories can vary greatly over depth (Bak et al., 2005; Hughes & Jackson, 1985; Johnston et al., 2021; Smith et al., 2016). Nonetheless, long-term monitoring efforts at mesophotic depths remain extremely rare, and published long-term datasets from tropical mesophotic coral reefs in the Indo-Pacific appear to be lacking entirely (Bongaerts et al., 2019).

Here, we present the results from permanent quadrats established concurrently at shallow (10 m) and upper mesophotic depths (40 m) in the Great Barrier Reef (GBR) and Western Coral Sea (WCS) (Fig. 1). During the 4-5 years of this study (from 2012 to 2017), several major disturbances (tropical cyclones and the 2016 mass bleaching) affected the communities across the seven monitoring sites, providing insight into the differential cumulative impact on shallow and mesophotic benthic communities. Using point-based tracking, we compare the trajectories of scleractinian cover and overall benthic community structure at shallow and mesophotic depths. Our results highlight different shallow and mesophotic scleractinian cover trajectories and reveal initial insights on the poorly understood dynamics between shallow and deep reef environments. Given the substantial area that mesophotic depths represent of the world’s coral reef habitat, we argue the urgency of incorporating MCEs in longitudinal monitoring efforts to understand and predict their role in a world increasingly impacted by climate change.

**Figure 1.**
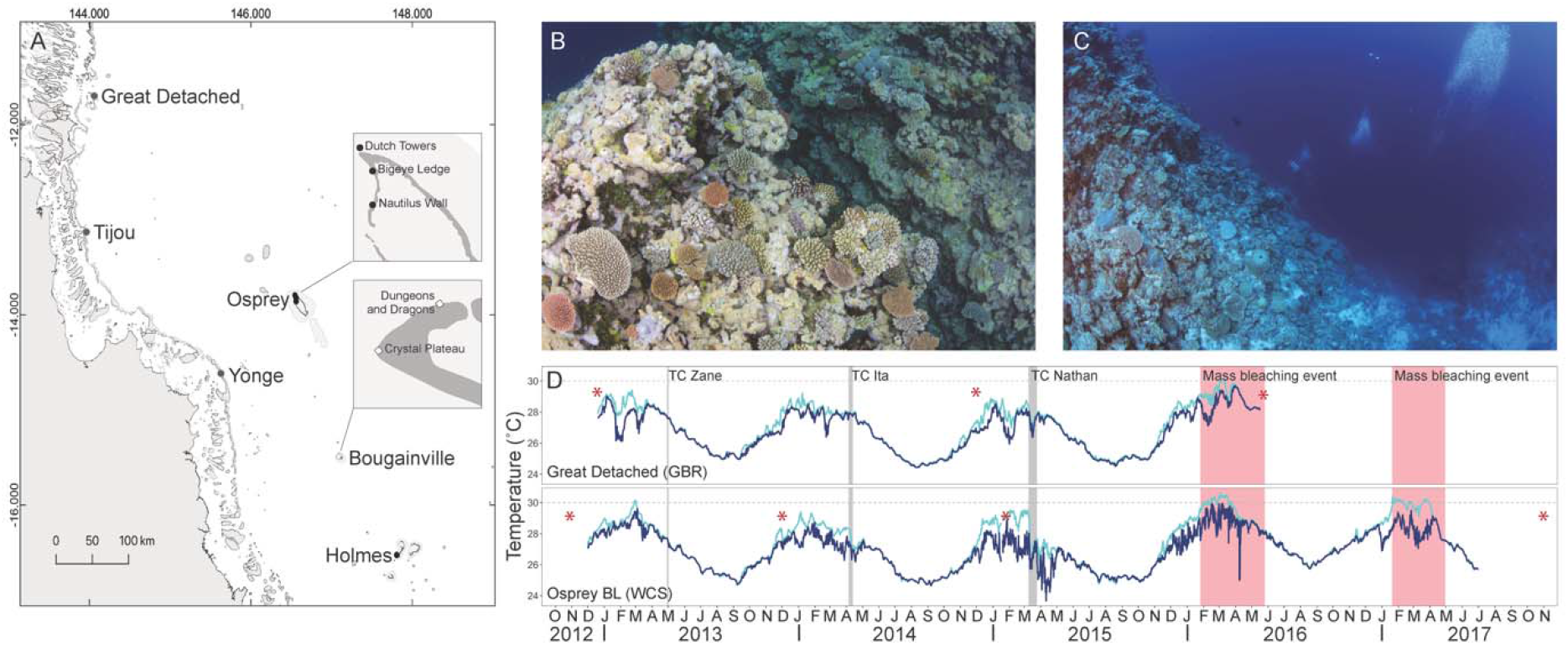
Locations and temperature profiles of the monitoring sites. (A) Map detailing the locations of the sampling sites. Great Barrier Reef (GBR) sites are indicated as grey dots, Western Coral Sea (WCS) reef sites with black dots. (B) Example of a shallow reef habitat (10 m) from Osprey DT. (C) Example of a mesophotic reef habitat (40 m) from Osprey DT. (D) Daily mean temperature profiles from one of the Great Barrier Reef (Great Detached) and Western Coral Sea (Osprey BL) locations between 2012 and 2017. The timeline also indicated the occurrence of three Tropical Cyclones (TCs) in grey and two mass bleaching events in red. Sampling dates are indicated with red asterisks. Daily mean temperature profiles of all the sampling sites in Fig. S1.

## Methods

### Establishment of permanent quadrats and image collection

Permanent quadrats were established in 2012 as part of the XL Catlin Seaview Survey at three locations in the far northern Great Barrier Reef (Great Detached Reef, Tijou Reef, and Yonge Reef) and two locations in the adjacent Western Coral Sea (Osprey Reef and Holmes Reef) (Fig. 1). The establishment and monitoring of permanent quadrats were conducted under permits 018-CZRS-1207626-01, 018-RRRW-131031-01 (Australian Department of the Environment), G12/35281.1, and G14/37294.1 (Great Barrier Reef Marine Park Authority). Monitoring was conducted following a methodology similar to that described by Bak & Nieuwland (1995). Three permanent quadrats (3 m × 3 m of electrical conduit PVC) were secured to the reef at both 10 m (shallow) and 40 m (upper mesophotic) depth using heavy-duty cable ties. At Osprey Reef, monitoring quadrats were established at three different sites, Osprey Dutch Towers (DT), Osprey Bigeye Ledge (BL), and Osprey Nautilus Wall (NW) (~4 km apart). Photographs were taken using a Canon EOS 5DmIII (with a 15 mm fisheye lens and a 1.4× teleconverter) in an Aquatica underwater housing and a Canon PowerShot G12 (standard lens at widest setting) in a Patima underwater housing, each fitted with two Sea & Sea YS-D1 underwater strobes. Overview (capturing the entire quadrat) and close-up (capturing ~0.5–1 m^2^) photographs were taken for each quadrat with both camera systems. Revisits were largely determined by opportunities for sea time and further limited by weather conditions but were preferably conducted in austral summer.

Since the initial survey of 2012, there were three tropical cyclones (TCs) affecting the study areas (Fig. 1D, Fig. S1): TC Zane from 29 April – 2 May 2013 (Category 1), TC Ita from 5–13 April 2014 (Category 5), TC Nathan from 9–25 March 2015 (Category 4). In addition, two large-scale bleaching events occurred in the late summers of 2016 and 2017. TC Zane, and possibly TC Ita for Yonge Reef, resulted in damage to shallow water quadrats at Holmes, Osprey NW, and Yonge Reef (seven out of nine quadrats were affected), but quadrat outlines could be reconstructed from the original photograph except for one quadrat at Yonge Reef, which was excluded from the dataset. Two locations at the Coral Sea, Bougainville Dungeons and Dragons (DD) and Bougainville Crystal Plateau (CP), were established in 2017 and only included for comparison (with no disturbance history). Map of locations (Fig. 1) was generated with the Group Layer “GBRMPA features” (Great Barrier Reef Marine Park Authority (GBRMPA), Copyright Commonwealth of Australia (2007)) and “Great Barrier Reef and Coral Sea Bathymetry” data set.

### Benthic community composition over time

The benthic cover was annotated using the CoralNet software (https://coralnet.ucsd.edu) (Beijbom et al., 2012) following a similar approach as de Bakker et al. (2017) identifying benthic categories intersected in a 225-point grid projected onto the overview quadrat photo (Fig. S2-S3). The grid system allowed for tracking of individual benthic points over time, with the original 2012 overview photo used as a reference to compensate for variation in height/angle of photographing between years. The points were reidentified in overview photos of subsequent years, using close-up imagery to facilitate the identification and resolve ambiguities. Individual points were categorized as ‘scleractinian corals’, ‘octocorals’, ‘crustose coralline algae (CCA)’, *‘Halimeda’*, ‘macroalgae’ (other than *Halimeda*), ‘turf algae’, ‘sponges’, *‘Millepora’*, ‘cyanobacteria’ and ‘unconsolidated substrate’ (as in de Bakker et al. (2017); González-Rivero et al. (2014)). *‘Halimeda’* was included as a separate category due to their major contribution to mesophotic benthic communities (Spalding et al., 2019). The groups ‘scleractinian corals’ and ‘octocorals’ were further categorized based on broad family/morphological classification (González-Rivero et al., 2014), to provide additional detail to benthic composition (Table S1). Points showing severely (i.e., completely white) bleached tissue of ‘scleractinian corals’ were separately classified. Other less abundant groups (e.g., tunicates, zoanthids, and anemones) were categorized as ‘other’. Points were categorized as ‘unidentifiable’ when identification was not possible (e.g., due to shading or camera angle).

### Statistical analyses

Time and space differences in the overall benthic community and scleractinian coral cover (absolute and relative change) were evaluated with a permutational multivariate analysis of variance (PERMANOVA+ (Anderson et al., 2008), Type III sum of squares and 9,999 permutations) on Bray-Curtis (community structure) and Euclidean (coral coverage) distances. Scleractinian coral cover was analyzed including bleached tissue unless otherwise indicated. The data were transformed (fourth root), standardized, and analyzed under a hierarchical design; *Region* (fixed), *Site* (random, nested in *Region*), *Depth* (fixed and orthogonal to *Site*), *quadrat* (random and nested in the interaction *Site×Depth*), and *Year* (fixed). The highest interaction of the model was excluded (*Quadrat(Site(Region)×Depth)×Year*). Correlations between univariate variables were evaluated using the R vegan package (https://cran.r-project.org/web/packages/vegan/index.html).

## Results and Discussion

Over the study period, our monitoring sites experienced two major cyclones and a mass bleaching event (Fig. 1D), which resulted in a significant reduction in coral cover (2012 vs 2017) (Fig. 2A; Fig. S4A, Tables S2-S3). Most of the change can be attributed to the drop in scleractinian coral cover at shallow depths (10 m) between the last two time points, after the combined impacts of TC Nathan and the 2016/2017 mass bleaching event (Fig. 2A). This corroborates impact patterns observed by other shallow reef surveys conducted in this region (Gonzalez-Rivero et al., 2017; Hughes et al., 2017). The impacts of TCs Zane (2013) and Ita (2014) were also apparent through extensive physical damage to the shallow quadrat PVC structures at Osprey NW, Holmes, and Yonge. In contrast, at mesophotic depths (40 m), no significant change in absolute scleractinian coral cover was observed (Fig. 2A) despite their direct adjacency to the shallow-water quadrats.

**Figure 2.**
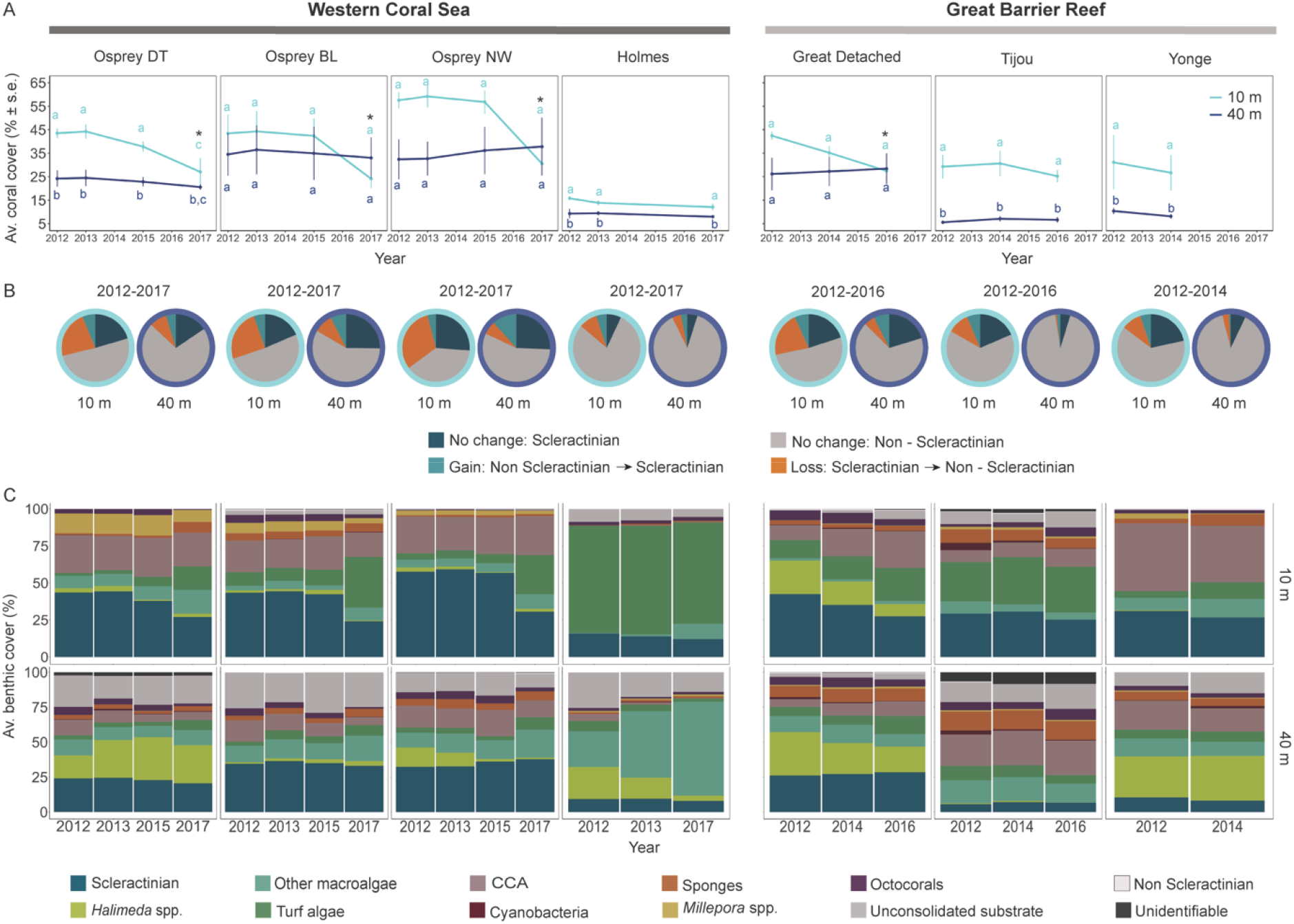
Changes in scleractinian coral cover and benthic community structure across time. (A) Scleractinian coral cover over time at shallow (10 m) and mesophotic (40 m) depths. Letters indicate significant differences between depths and the interaction *DepthxYear* (*p*<0.05, Table S4), and asterisks overall differences between 2012 and 2017 (*p*<0.05, Table S5). Error bars indicate standard error across replicate quadrats (s.e.). (B) Gains and losses in scleractinian cover based on point-based tracking (225 points per quadrat) between 2012 and 2016/2017 (raw data in Fig. S2-S3). (C) Changes in community structure at shallow (10 m) and mesophotic (40 m) depths. “Other macroalgae” represent all macroalgae species excluding *Halimeda* spp (listed separately). “CCA” stands for crustose coralline algae.

Given that absolute loss in coral cover is known to positively correlate with initial cover (Osborne et al., 2011), we also looked at the proportional change between 2012 and 2017. This was also significantly higher at shallow depths (~38% versus ~2% at shallow and mesophotic depths respectively; Fig. S4B, Tables S6). In fact, while seven sites showed a relative decrease in shallow scleractinian coral cover in the final year, three out of seven sites showed a relative increase in mesophotic cover (Fig. 2A, Fig. S4B). At shallow depths, the relative change in scleractinian cover was also negatively correlated with initial cover (Fig. 3A), which is not unexpected as higher cover may relate to larger coral structures that are more susceptible to storm damage (Madin et al., 2014). However, such a correlation was not observed at mesophotic depths, highlighting that the lack of decline at these depths was not merely due to lower coral cover (Fig. 3A) (as the relative change was similar for high-cover sites, e.g., Osprey BL and NW).

**Figure 3.**
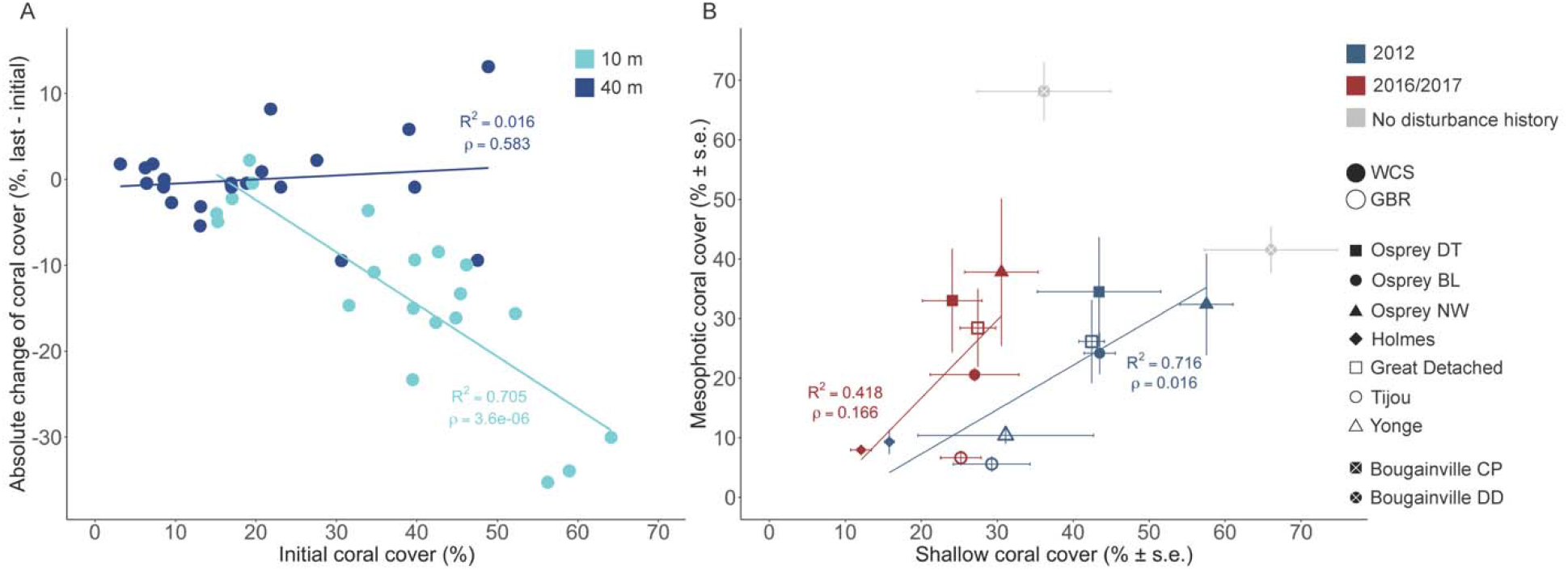
Changes and relationships between scleractinian coral cover at shallow (10 m) and mesophotic (40 m) depths. (A) Correlation between the initial scleractinian coral cover (2012) and the absolute change over time. (B) Correlation between shallow and mesophotic coral cover at the beginning of the study period (2012) and at the end, after several disturbances (2016/2017). Bougainville DD and Bougainville CP were only surveyed at the last time point (2017). Error bars indicate standard error across replicate quadrats (s.e.).

Initially, there was a clear positive correlation between shallow and mesophotic scleractinian cover across locations, despite scleractinian coral cover varying greatly across sites and depths (Fig. 3B). Although such shallow-mesophotic correlations are rarely explored, it is perhaps not surprising given that the mesophotic communities occur as direct extensions of shallow reef habitat, and their development may be determined by similar environmental requirements as their shallow-water counterparts (Gress et al., 2018). For example, hard substrate availability ultimately determines the potential for scleractinian coral cover and is strongly determined by local geomorphological features that can extend from shallow to mesophotic depths (Sherman et al., 2019). Similarly, certain chronic pressures such as sedimentation, ocean acidification, pollution, disease outbreaks, and invasive species are expected to impact coral communities across broad depth ranges (Bak et al., 2005; Smith et al., 2019), further contributing to this shallow-mesophotic correlation. After the disturbances (2016/2017), this correlation broke down as the acute disturbances predominantly impacted shallow scleractinian cover, with mesophotic scleractinian cover remaining at similar levels (Fig. 3B). Although the pre-disturbance correlation indicates a clear relationship between shallow-mesophotic coral cover, the post-disturbance shift highlights how the differential impact of disturbances over depth can lead to diverging benthic trajectories.

Changes in coral cover represent a net balance of gains (through e.g. coral growth and recruitment) and losses (through e.g. mortality and displacement) (Brito-Millán et al., 2019; Hughes & Jackson, 1985). Point-based tracking of the benthic community within our permanent quadrats revealed that the relatively consistent cover at mesophotic depths is driven by near-identical gains and losses (Fig. 2B, 6 ± 1.1 % and 5.7 ± 0.9 %, respectively, excluding Yonge) resulting in a net change close to zero. Although the hypothesized stability of MCEs is often colloquially interpreted as these ecosystems being more static (Smith et al., 2019), early quadrat-based observations already pointed towards their potentially dynamic nature (Bak et al., 2005; Liddell & Avery, 2000). Visual observations of the quadrats support this notion, with roughly a third of scleractinian coral colonies suffering partial mortality and many examples of complete colony displacements (particularly in tabular *Acropora* colonies). At the same time, considerable growth rates were observed for both tabular and plating corals at mesophotic depths, with anecdotal measurements including radial extension rates of ~2-4 cm/year in *Acropora, Montipora, Mycedium*, and *Pachyseris*. We also observed substantial shifts in community structure, with Acroporidae and Poritidae decreasing in coverage over time at mesophotic depths (7.4% and 7.7%, respectively), while all other families, Faviidae-Mussidae, Pocilloporidae, Merulinidae, Agariciidae, and Fungiidae, increased in coverage between 5% to 37% (Fig. S5). Regarding the morphological composition of the mesophotic reef, branching and plating morphologies increased 51% and 19% respectively (to their initial coverage), whereas massive, encrusting, and corymbose morphologies decreased between 2 and 24% (Fig. S6).

In terms of the overall benthic structure, Scleractinia, *Millepora* (fire coral), crustose coralline algae, and unconsolidated substrate drove the overall differences between depths (Fig. S7-S8). Shallow reef communities showed increases in turf algae, and decreases in coral cover, similar to trajectories observed in many disturbance-affected reefs (Jackson et al., 2014). Macroalgae (primarily *Halimeda* and fleshy algae) dominated and were major drivers of change in mesophotic communities, although coverage and trajectories were highly variable across sites (Fig. 2C). For example, at Osprey DT, *Halimeda* doubled its coverage in a single year (2012-2013), whereas in Osprey NW and Holmes was almost extirpated by 2017. At Holmes, the disappearance of *Halimeda* coincided with an outbreak of *Microdictyon* (from 25% to 67%), which colonizes mesophotic reefs by forming large monospecific mats (Kahng et al., 2017; Spalding et al., 2019). Macroalgae dominance in mesophotic reefs have been previously reported in the Western Atlantic and Indo-Pacific, playing a fundamental role in the geomorphology of mesophotic habitats by consolidating and modifying the complexity of the substrate (Spalding et al., 2019). Observed changes in macroalgae abundance can likely be attributed to seasonal fluctuations in temperature and nutrients related to upwelling intensity, to which macroalgae species can be responsive (Spalding et al., 2019).

Overall, these results demonstrate the importance of concurrent monitoring reefs at shallow and mesophotic depths and yielded some critical insights: (1) The initial correlation between shallow and mesophotic cover highlighted the connection between benthic communities at different depths, a fundamental observation that warrants further investigation to understand the underlying drivers of this relationship. (2) The observed breakdown of the correlation following disturbances led to divergent trajectories in directly adjacent communities (shallow and mesophotic), challenging us to question whether and how such relationships naturally rebound. (3) Lastly, in contrast to their conventional view as slow-growing and “static”, the observed dynamic nature of scleractinian coral communities at mesophotic depths highlights the rapid changes these communities can undergo independent from shallow communities. Although monitoring efforts at mesophotic depths are inherently more costly and logistically complicated, they are gradually becoming more accessible through advances in underwater (Armstrong et al., 2019; Pyle, 2019) and imaging technologies (e.g. Bongaerts et al. (2021); Ferrari et al. (2016)). We consider that such efforts need to be prioritized given the major spatial extent of these communities and their role in the overall coral reef ecosystem. Only through sustained observations at these depths will we get the insights needed to understand the fundamental ecological processes that govern these ecosystems, and ensure that we maximize our ability to protect them into the future.

## Acknowledgements

We thank David Whillas, Kyra Hay and Paul Muir for their help with the initial establishment and rephotographing of the permanent quadrats. For field support, we would like to thank Jaap Barendrecht, Linda Tonk, David Aguirre, Pedro Frade, Manuel Gonzalez-Rivero, David Harris, Susie Green, Erin McFadden, Steve Lindfield, Sara Naylor, as well as Underwater Earth, The Ocean Agency, and crews from Reef Connections, Mike Ball Dive Expeditions, SY Ethereal, and the Waitt Foundation. We thank Juan J. Cruz-Motta for comments regarding data analysis. This project was supported (in chronological order) by the XL Catlin Seaview Survey (2012–2014; funded by the XL Catlin Group in partnership with Underwater Earth and The University of Queensland), an Australian Research Council Discovery Early Career Researcher Award (2016–2018; DE160101433), and the Hope for Reefs Initiative at the California Academy of Sciences. Substantial sea time was generously provided by the Waitt Foundation and the Joy Foundation.

## Author contributions

PB, NE and OHG conceived and designed the study. PB and NE conducted the fieldwork. FS conducted the image classification. AHA, FS and PB conducted and contributed to the analyses. AHA, PB and FS wrote the manuscript and all the authors contributed to the edits.

## Data accessibility statement

Annotation and temperature data will be available through https://github.com/pimbongaerts/monitoring.

## Notes

https://github.com/pimbongaerts/monitoring

### Competing Interest Statement

The authors have declared no competing interest.

