## Supplementary material for "Hidden in the deep: distinct benthic trajectories call for monitoring of mesophotic reefs": Fig. S

### 1.1 Supplementary Figures

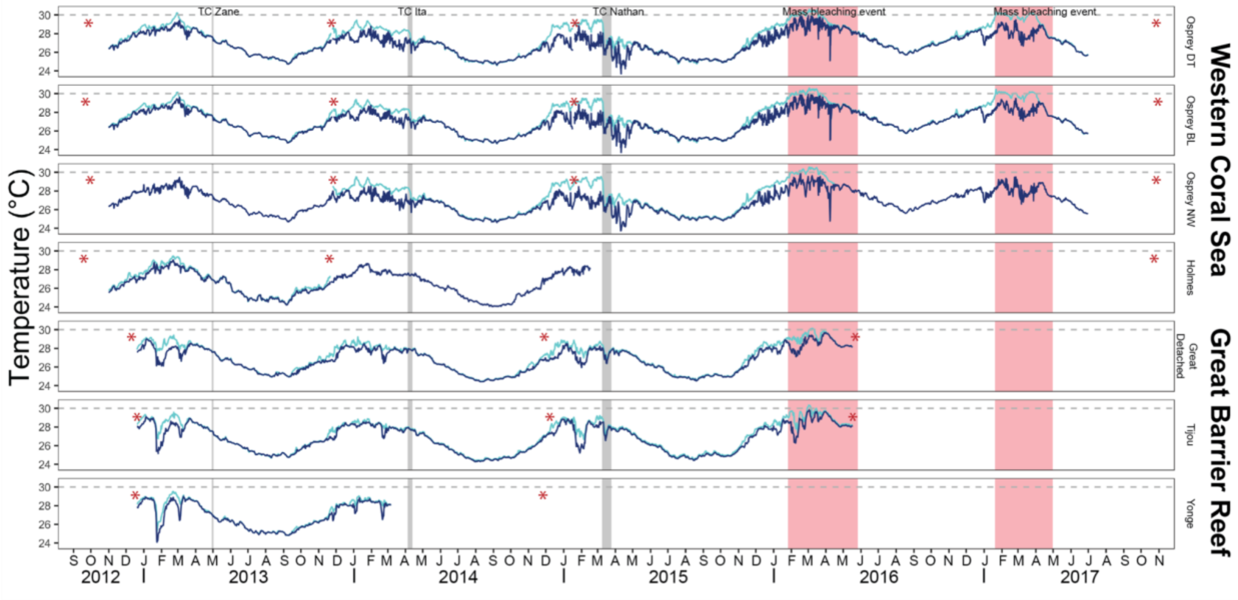

**Figure S1.** Daily mean temperature profiles in locations of the Western Coral Sea (Osprey DT, Osprey BL, Osprey NW, and Holmes) and the Great Barrier Reef (Great Detached, Tijou, and Yonge) between 2012 and 2017. The timeline also indicated three Tropical Cyclones (TCs) in grey and two mass bleaching events in red. Sampling dates are indicated with red asterisks.

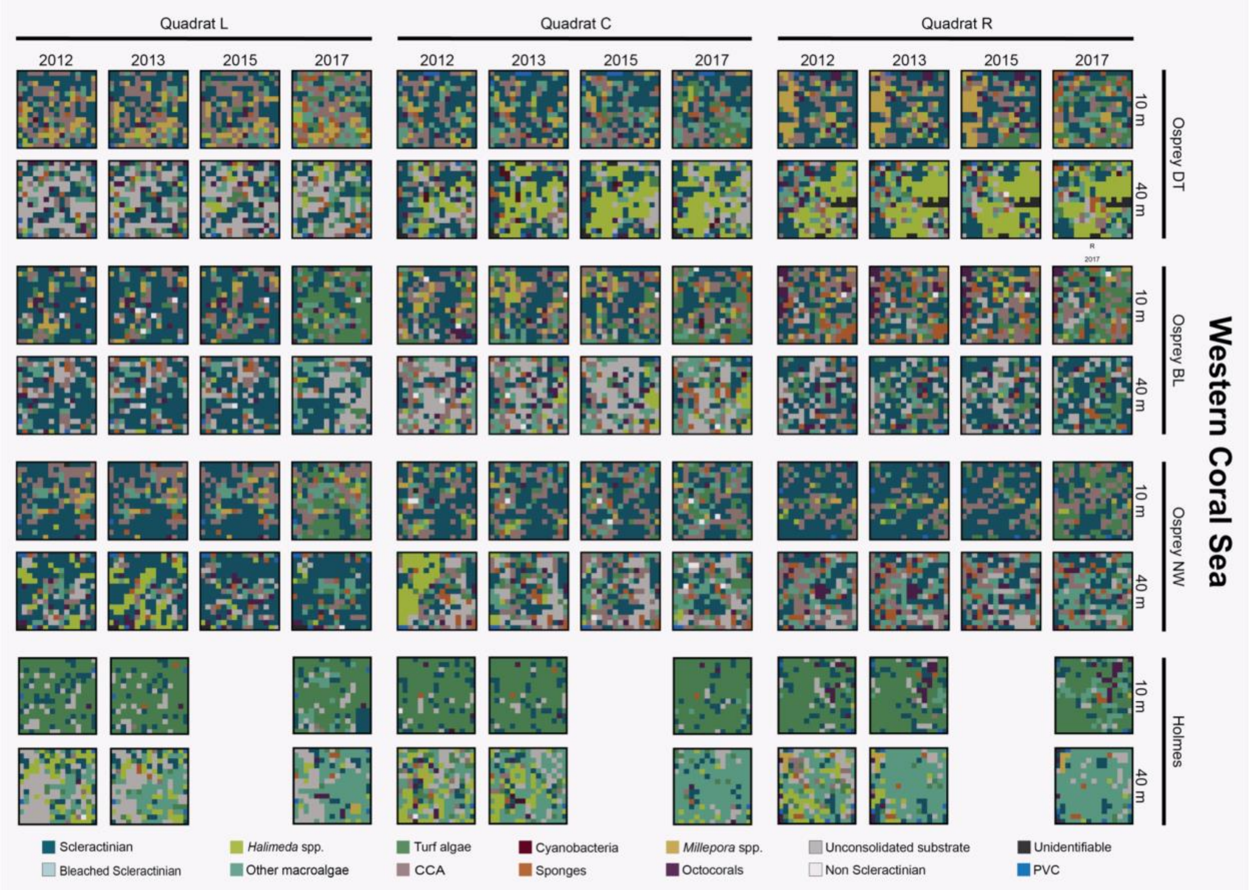

**Figure S2.** Benthic cover in individual quadrats of the Western Coral Sea (raw data). Each box represents 3 m x 3 m quadrats, and tiles inside are the benthic categories identified in each of the 225-point grids projected onto the overview quadrat photo. Empty spaces indicate quadrats not sampled at that time (see details in the Methods section). “Other macroalgae” represent all macroalgae species excluding *Halimeda* spp (listed separately). “CCA” stands for crustose coralline algae.

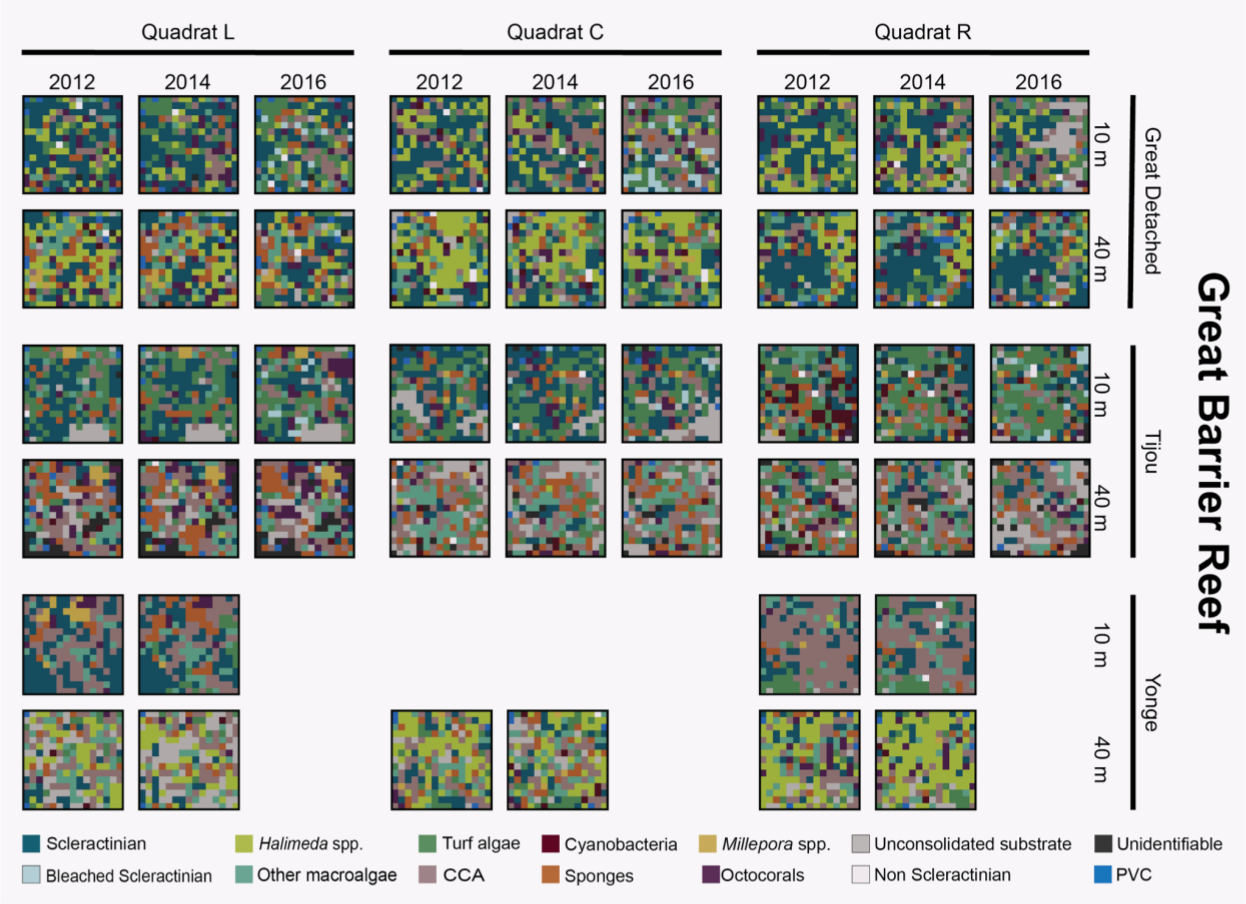

**Figure S3.** Benthic cover in individual quadrats of the Great Barrier Reef (raw data). Each box represents 3 m x 3 m quadrats, and tiles inside are the benthic categories identified in each of the 225-point grids projected onto the overview quadrat photo. Empty spaces indicate quadrats not sampled at that time (see details in the Methods section). “Other macroalgae” represent all macroalgae species excluding *Halimeda* spp (listed separately). “CCA” stands for crustose coralline algae.

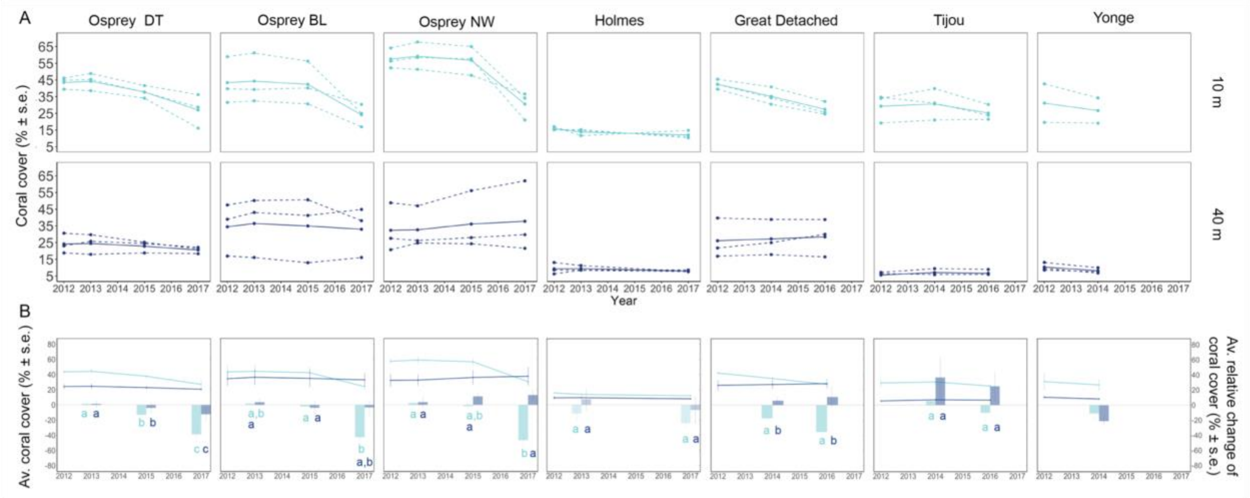

**Figure S4.** Coral coverage changes over time. (A) Coral coverage over time in individual quadrats. (B) Relative change of coral cover over time (based on 2012 coverage). Trajectories are indicated with lines (left y-axis) and the relative change as bars (right y-axis). Letters indicate significant differences (Tables S6). For significance in the trajectories, see Fig. 2 in the main text and Tables S4-S5. Error bars indicate standard error across replicate quadrats (s.e.).

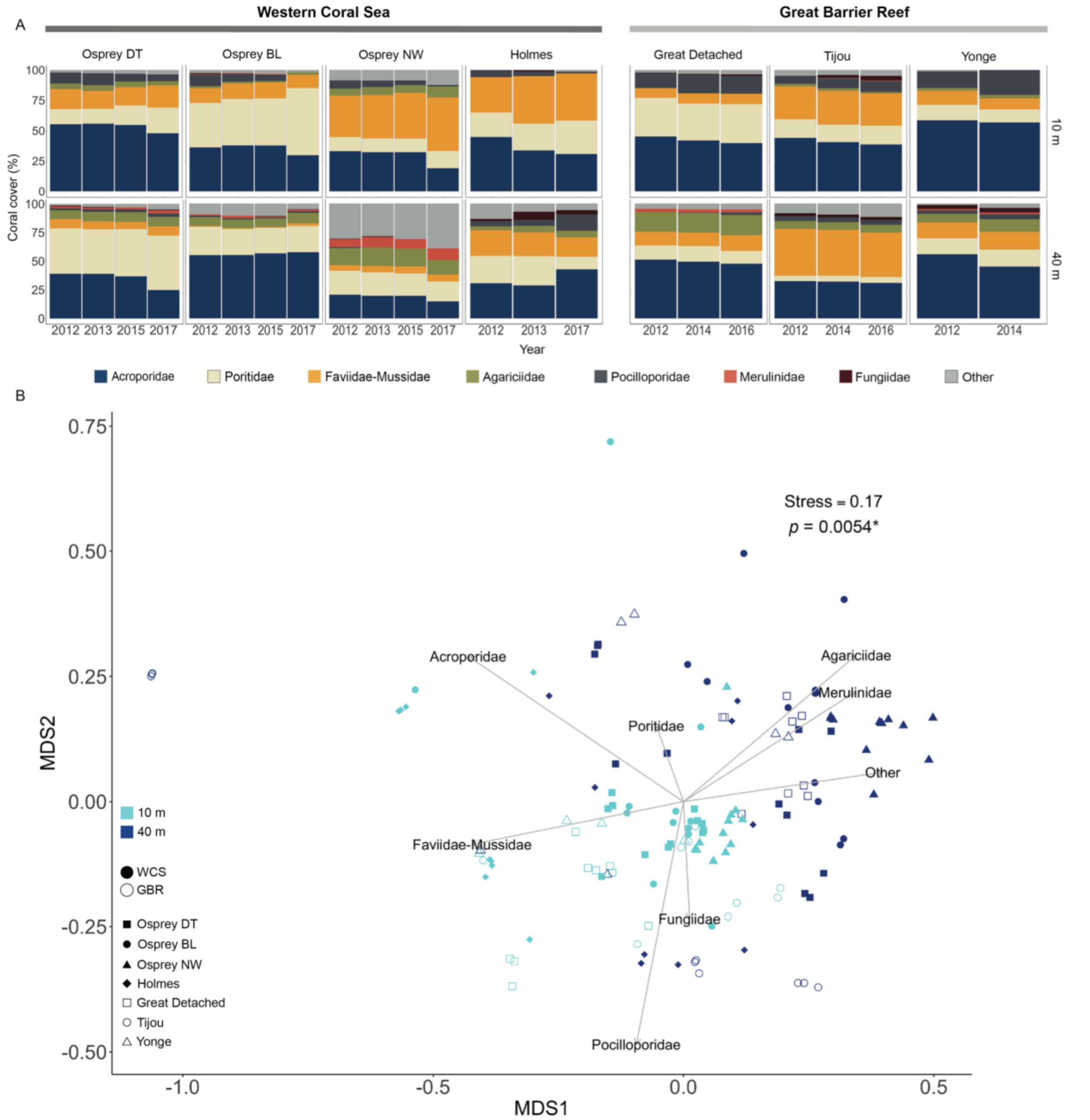

**Figure S5.** Structure of scleractinian coral community (A) and non-Multidimensional Scaling (B) by families. Points in the nMDS represent a time point for each quadrat. *P*-value with an asterisk (*) refers to a significant difference between depths (see Table S7).

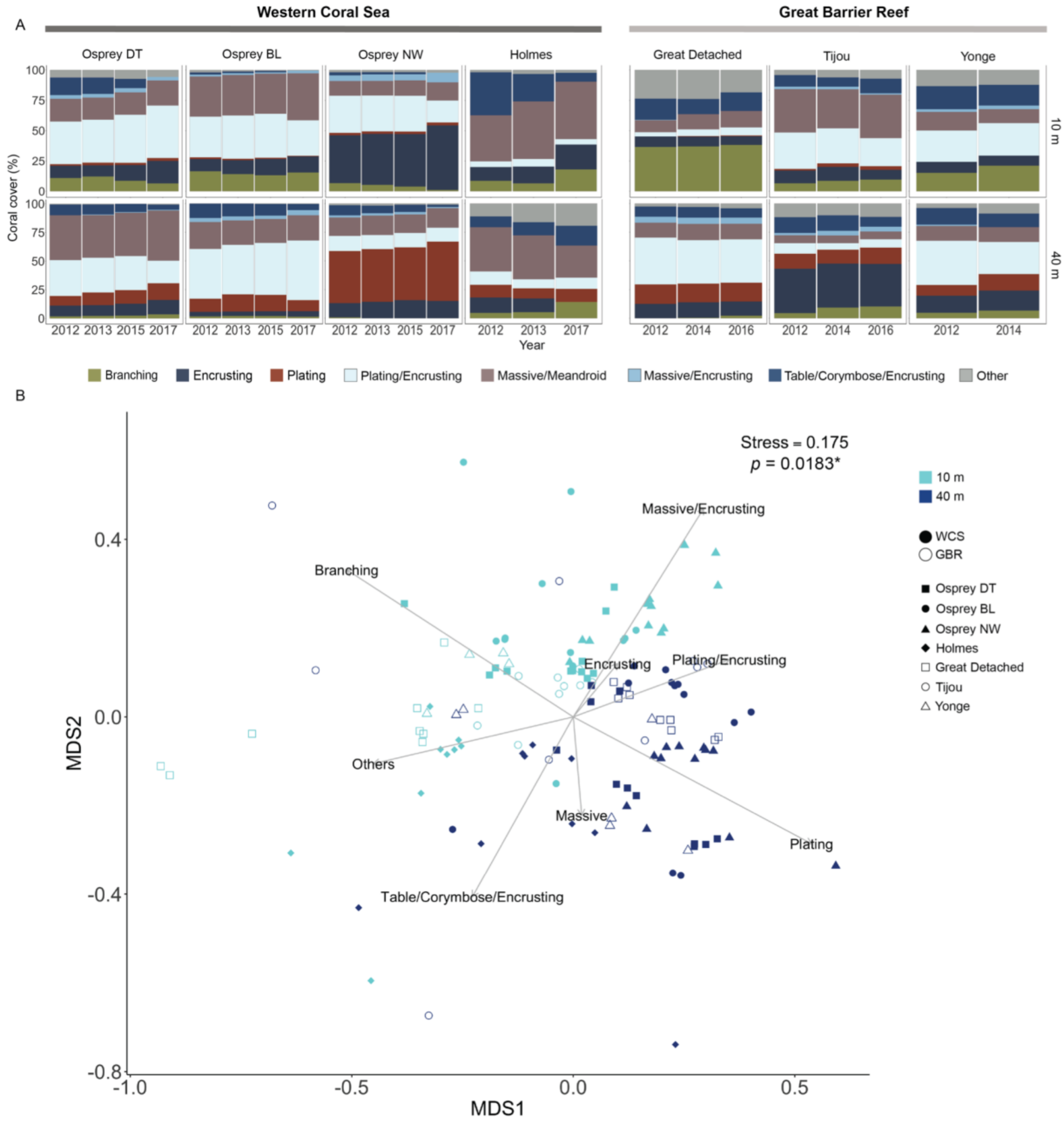

**Figure S6.** Structure of scleractinian coral community (A) and non-Multidimensional Scaling (B) by morphologies. Points in the nMDS represent a time point for each quadrat. *P*-value with an asterisk (*) refers to a significant difference between depths (see Table S8).

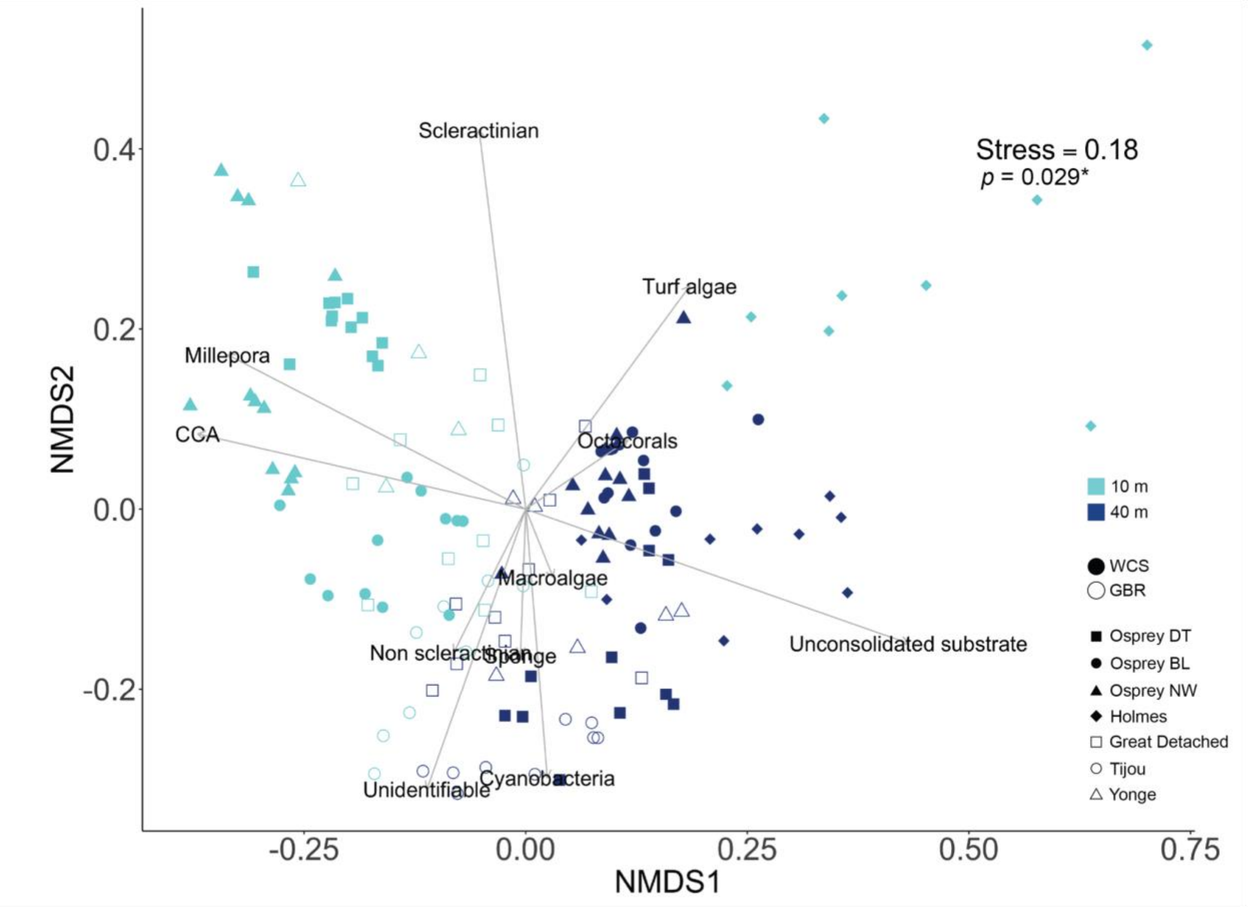

**Figure S7.** Non-Multidimensional Scaling (nMDS) with vector overlays of major benthic groups. Points in the nMDS represent a time point for each quadrat. "Macroalgae" groups all macroalgae species, including *Halimeda* spp. *P*-value with an asterisk (*) refers to a significant difference between depths (see Table S9).

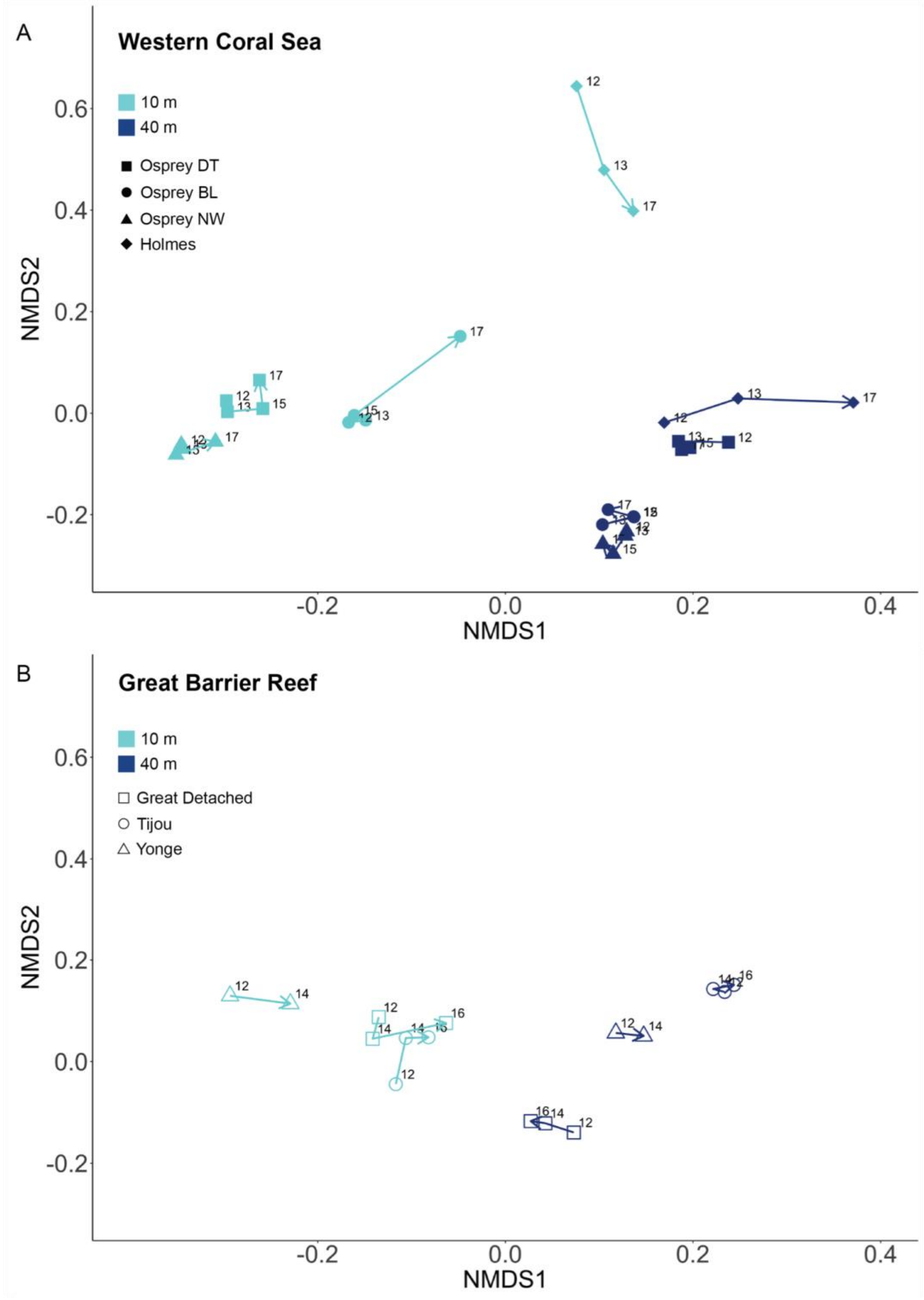

**Figure S8.** Temporal trajectories of benthic community structure in the Western Coral Sea (A) and the Great Barrier Reef (B). Points in the nMDS represent the centroid of the quadrats per site, year, and depth. Numbers indicate the last two digits of the year and arrows the trajectory from 2012 to 2016/2017.

### 1.2 Supplementary Tables

**Table S1. Sub-categories for scleractinian corals and octocorals.**

| Benthic group | Family or genus (italics) | Morphology |
|---|---|---|
| Scleractinian corals | Acroporidae | Branching |
| | | Encrusting |
| | | Table/corymbose/encrusting |
| | | Plate/encrusting |
| | | Other |
| | Faviidae-Mussidae | Encrusting |
| | | Massive/meandroid |
| | | Plating |
| | | Other |
| | Fungiidae | Plating |
| | | Other |
| | Poritidae | Branching |
| | | Encrusting |
| | | Massive |
| | | Plating |
| | | Other |
| | *Seriatopora* | Branching |
| | *Stylophora* | Branching |
| | *Pocillopora* | Branching |
| | *Mycedium* | Plating |
| | *Leptoseris* | Encrusting |
| | | Plating |
| | | Plating/encrusting |
| | *Pachyseris* | Plating |
| | *Pavona* | Massive/encrusting |
| | Other hard coral | Encrusting |
| | | Massive/meandroid |
| | | Plating |
| | | Other |

| | |
|---|---|
| Octocorals | Alcyoniidae |
| | Sarcophyton |
| | Other Soft Coral |
| | Gorgonians |

**Table S2. Permutational analysis of variance (univariate PERMANOVA) on scleractinian coral coverage.** Test based on Euclidean distances on fourth root transformed and standardized data. Significant differences are indicated in bold. P(perm): *P*-value based on permutations, U. perms: Unique permutations, P(MC): Monte Carlo *P*- value, ECV(%): Estimated components of variation.

| Source | df | SS | MS | Pseudo-F | P(perm) | U. perms | P(MC) | ECV (%) |
|---|---|---|---|---|---|---|---|---|
| Region | 1 | 105.35 | 105.35 | 7.9615 | **0.0337** | 9865 | **0.0309** | 16.2 |
| Depth | 1 | 400.02 | 400.02 | 10.14 | **0.0208** | 9838 | **0.0212** | 20.5 |
| Year | 5 | 52.575 | 10.515 | 4.8731 | **0.0141** | 9952 | **0.0162** | 5.0 |
| Site(Region) | 5 | 202.04 | 40.408 | 2.7172 | **0.0442** | 9947 | **0.041** | 8.7 |
| RegionxDepth | 1 | 3.8554 | 3.8554 | 0.2301 | 0.6579 | 9861 | 0.6543 | 0.0 |
| RegionxYear** | 0 | 0 | | No test | | | | 0.0 |
| DepthxYear | 5 | 60.425 | 12.085 | 10.537 | **0.0007** | 9946 | **0.0007** | 8.1 |
| Site(Region)xDepth | 5 | 260.03 | 52.006 | 3.4971 | **0.0173** | 9956 | **0.014** | 14.9 |
| Site(Region)xYear** | 11 | 23.761 | 2.1601 | 1.79 | 0.0715 | 9933 | 0.0789 | 3.0 |
| RegionxDepthxYear** | 0 | 0 | | No test | | | | 0.0 |
| Quadrat(Site(Region)xDepth) | 27 | 416.9 | 15.441 | 12.795 | **0.0001** | 9900 | **0.0001** | 15.5 |
| Site(Region)xDepthxYear** | 11 | 12.614 | 1.1467 | 0.95028 | 0.4977 | 9917 | 0.4921 | 0.0 |
| Residual | 63 | 76.024 | 1.2067 | | | | | 8.2 |
| Total | 135 | 1969.2 | | | | | | |

**Table S3. Pairwise comparisons from permutational multivariate analysis of variance (univariate PERMANOVA) for the factors *Site* and *Year* on scleractinian coral cover data.** Test based on Euclidean distances on fourth root transformed and standardized data. Significant differences are indicated in bold. P(perm): *P*-value based on permutations, U. perms: Unique permutations, P(MC): Monte Carlo *P*- value, ECV(%): Estimated components of variation.

*Site(Region)*

| **Region** | **Groups** | **t** | **P(perm)** | **U. perms** | **P(MC)** |
|---|---|---|---|---|---|
| Coral Sea | HOLME, OSPBL | 0.0058647 | 0.9899 | 9249 | 0.9963 |
| | HOLME, OSPDT | 0.14973 | 0.8847 | 9250 | 0.8865 |
| | HOLME, OSPNW | 1.5434 | 0.174 | 9309 | 0.1574 |
| | OSPBL, OSPDT | 0.16253 | 0.8631 | 8705 | 0.8748 |
| | OSPBL, OSPNW | 1.5413 | 0.1615 | 8794 | 0.1599 |
| | OSPDT, OSPNW | 2.1795 | 0.063 | 8811 | 0.0642 |
| Great Barrier Reef | GRDET, TIJOU | 2.9039 | **0.0166** | 8779 | **0.0233** |
| | GRDET, YONGE | 0.4461 | 0.6799 | 8851 | 0.676 |
| | TIJOU, YONGE | 2.5125 | **0.0306** | 8869 | **0.0381** |

*Year*

| **Groups** | **t** | **P(perm)** | **U. perms** | **P(MC)** |
|---|---|---|---|---|
| 2012, 2013 | 1.7516 | 0.1908 | 5393 | 0.1835 |
| 2012, 2014 | 1.0501 | 0.3969 | 1119 | 0.3987 |
| 2012, 2015 | 0.34666 | 0.7795 | 427 | 0.7618 |
| 2012, 2016 | 0.33821 | 0.8732 | 21 | 0.7929 |
| 2012, 2017 | 8.2401 | **0.0031** | 5434 | **0.004** |
| 2013, 2015 | 0.88486 | 0.4785 | 212 | 0.4798 |
| 2013, 2017 | 4.0691 | 0.0521 | 425 | **0.0278** |
| 2014, 2016 | 0.35472 | 0.6449 | 15 | 0.7784 |
| 2015, 2017 | 3.1169 | 0.1311 | 212 | 0.0935 |

**Table S4. Permutational analysis of variance (univariate PERMANOVA) on scleractinian coral coverage over time by site and pairwise comparisons for the interactions *DepthxYear* and *Year*.** Test based on Euclidean distances on fourth root transformed and standardized data. Significant differences are indicated in bold. P(perm): *P*-value based on permutations, U. perms: Unique permutations, P(MC): Monte Carlo *P*- value, ECV(%): Estimated components of variation.

#### *A) Coral Sea: Osprey – Dutch Towers*

| Source | df | SS | MS | Pseudo-F | P(perm) | U. perms | P(MC) | ECV (%) |
|---|---|---|---|---|---|---|---|---|
| Depth | 1 | 116.38 | 116.38 | 20.971 | 0.1018 | 10 | **0.0105** | 49.7 |
| Year | 3 | 10.916 | 3.6386 | 15.468 | **0.0007** | 9967 | **0.0006** | 12.3 |
| Quadrat(Depth) | 4 | 22.199 | 5.5498 | 23.592 | **0.0001** | 9955 | **0.0001** | 18.9 |
| DepthxYear | 3 | 4.8708 | 1.6236 | 6.9019 | **0.0071** | 9969 | **0.0059** | 11.1 |
| Residual | 12 | 2.8229 | 0.23524 | | | | | 7.9 |
| Total | 23 | 157.19 | | | | | | |

*DepthxYear by Depth*

| Depth | Groups | t | P(perm) | U. perms | P(MC) |
|---|---|---|---|---|---|
| 10 m | 2012, 2013 | 0.6684 | 0.6037 | 38 | 0.5705 |
| | 2012, 2015 | 0.43673 | 0.597 | 38 | 0.7055 |
| | 2012, 2017 | 7.1426 | 0.1009 | 38 | **0.0182** |
| | 2013, 2015 | 0.93135 | 0.4267 | 38 | 0.4476 |
| | 2013, 2017 | 4.6215 | 0.0972 | 38 | **0.0437** |
| | 2015, 2017 | 4.5224 | 0.0962 | 38 | **0.045** |
| 40 m | 2012, 2013 | 1.7411 | 0.2788 | 38 | 0.2313 |
| | 2012, 2015 | 1.724 | 0.3196 | 38 | 0.2276 |
| | 2012, 2017 | 3.5205 | 0.1726 | 38 | 0.0723 |
| | 2013, 2015 | 1.0243 | 0.4641 | 38 | 0.4161 |
| | 2013, 2017 | 1.3821 | 0.3289 | 38 | 0.2966 |
| | 2015, 2017 | 1.0898 | 0.3135 | 38 | 0.3944 |

*DepthxYear by Year*

| Year | Groups | t | P(perm) | U. perms | P(MC) |
|---|---|---|---|---|---|
| 2012 | 10m, 40m | 4.7886 | 0.1038 | 10 | **0.0095** |
| 2013 | | 7.1615 | 0.1049 | 10 | **0.0027** |
| 2015 | | 4.6417 | 0.098 | 10 | **0.0092** |
| 2017 | | 2.2891 | 0.198 | 10 | 0.0817 |

#### ***B) Coral Sea: Osprey – Bigeye Ledge***

| Source | df | SS | MS | Pseudo-F | P(perm) | U. perms | P(MC) | ECV (%) |
|---|---|---|---|---|---|---|---|---|
| Depth | 1 | 12.685 | 12.685 | 0.39555 | 0.5979 | 10 | 0.5548 | 0.0 |
| Year | 3 | 9.7127 | 3.2376 | 4.737 | **0.0221** | 9962 | **0.0203** | 12.2 |
| Quadrat(Depth) | 4 | 128.28 | 32.069 | 46.921 | **0.0001** | 9951 | **0.0001** | 52.3 |
| DepthxYear | 3 | 12.386 | 4.1288 | 6.041 | **0.0058** | 9950 | **0.01** | 20.0 |
| Residual | 12 | 8.2016 | 0.68347 | | | | | 15.4 |
| Total | 23 | 171.26 | | | | | | |

*DepthxYear by Depth*

| Depth | Groups | t | P(perm) | U. perms | P(MC) |
|---|---|---|---|---|---|
| 10 m | 2012, 2013 | 0.0023525 | 1 | 37 | 0.9984 |
| | 2012, 2015 | 0.70436 | 0.6297 | 37 | 0.5545 |
| | 2012, 2017 | 3.1002 | 0.1686 | 49 | 0.0848 |
| | 2013, 2015 | 0.35185 | 1 | 38 | 0.7574 |
| | 2013, 2017 | 2.6531 | 0.1656 | 38 | 0.1166 |
| | 2015, 2017 | 2.8937 | 0.1679 | 38 | 0.1001 |
| 40 m | 2012, 2013 | 2.6997 | 0.1665 | 38 | 0.1167 |
| | 2012, 2015 | 3.1891 | 0.169 | 38 | 0.0897 |
| | 2012, 2017 | 0.69657 | 0.5668 | 38 | 0.551 |
| | 2013, 2015 | 1.4877 | 0.3517 | 38 | 0.2753 |
| | 2013, 2017 | 0.86667 | 0.5724 | 38 | 0.4771 |
| | 2015, 2017 | 0.46672 | 0.7332 | 38 | 0.6803 |

*DepthxYear by Year*

| Year | Groups | t | P(perm) | U. perms | P(MC) |
|---|---|---|---|---|---|
| 2012 | 10m, 40m | 0.54151 | 0.6967 | 10 | 0.6251 |
| 2013 | | 0.15354 | 0.806 | 10 | 0.8908 |
| 2015 | | 0.15145 | 1 | 10 | 0.8845 |
| 2017 | | 1.4863 | 0.303 | 10 | 0.209 |

*Year*

| Groups | t | P(perm) | U. perms | P(MC) |
|---|---|---|---|---|
| 2012, 2013 | 1.7575 | 0.1544 | 9577 | 0.1483 |
| 2012, 2015 | 0.88399 | 0.4276 | 9429 | 0.431 |
| 2012, 2017 | 3.0826 | **0.0384** | 9561 | **0.0401** |
| 2013, 2015 | 0.71335 | 0.502 | 9577 | 0.5203 |
| 2013, 2017 | 1.8155 | 0.1565 | 9612 | 0.1404 |
| 2015, 2017 | 2.2709 | 0.0866 | 9596 | 0.0873 |

***C) Coral Sea: Osprey – Nautilus Wall***

| Source | df | SS | MS | Pseudo-F | P(perm) | U. perms | P(MC) | ECV (%) |
|---|---|---|---|---|---|---|---|---|
| Depth | 1 | 90.371 | 90.371 | 2.848 | 0.1988 | 10 | 0.1708 | 25.0 |
| Year | 3 | 34.148 | 11.383 | 9.8537 | **0.0006** | 9959 | **0.0019** | 14.7 |
| Quadrat(Depth) | 4 | 126.93 | 31.732 | 27.47 | **0.0001** | 9964 | **0.0001** | 31.2 |
| DepthxYear | 3 | 23.769 | 7.9228 | 6.8587 | **0.005** | 9956 | **0.0078** | 17.0 |
| Residual | 12 | 13.862 | 1.1552 | | | | | 12.1 |
| Total | 23 | 289.08 | | | | | | |

*DepthxYear by Depth*

| Depth | Groups | t | P(perm) | U. perms | P(MC) |
|---|---|---|---|---|---|
| 10 m | 2012, 2013 | 1.3874 | 0.3504 | 38 | 0.307 |
| | 2012, 2015 | 1.5687 | 0.2537 | 38 | 0.2536 |
| | 2012, 2017 | 3.4868 | 0.1035 | 38 | 0.0707 |
| | 2013, 2015 | 1.4825 | 0.2848 | 38 | 0.2744 |
| | 2013, 2017 | 3.3931 | 0.1022 | 38 | 0.0749 |
| | 2015, 2017 | 3.1192 | 0.1649 | 38 | 0.0885 |
| 40 m | 2012, 2013 | 0.63428 | 0.552 | 38 | 0.5938 |
| | 2012, 2015 | 2.3241 | 0.2353 | 38 | 0.1429 |
| | 2012, 2017 | 0.23707 | 1 | 38 | 0.8365 |
| | 2013, 2015 | 1.3516 | 0.2787 | 38 | 0.3144 |
| | 2013, 2017 | 0.053971 | 1 | 38 | 0.9605 |
| | 2015, 2017 | 2.6289 | 0.2013 | 38 | 0.121 |

*DepthxYear by Year*

| Year | Groups | t | P(perm) | U. perms | P(MC) |
|---|---|---|---|---|---|
| 2012 | 10m, 40m | 2.1399 | 0.1023 | 10 | 0.0978 |
| 2013 | | 2.1482 | 0.1001 | 10 | 0.0945 |
| 2015 | | 1.6881 | 0.0985 | 10 | 0.1636 |
| 2017 | | 0.2459 | 0.9001 | 10 | 0.8192 |

*Year*

| Groups | t | P(perm) | U. perms | P(MC) |
|---|---|---|---|---|
| 2012, 2013 | 1.0186 | 0.3739 | 9567 | 0.3631 |
| 2012, 2015 | 2.4396 | 0.075 | 9634 | 0.0722 |
| 2012, 2017 | 3.2391 | **0.0359** | 9538 | **0.0352** |
| 2013, 2015 | 1.997 | 0.1199 | 9644 | 0.1136 |
| 2013, 2017 | 3.1188 | **0.039** | 9601 | **0.0379** |
| 2015, 2017 | 3.5363 | **0.0275** | 9588 | **0.0226** |

***D) Coral Sea: Osprey – Holmes***

| Source | df | SS | MS | Pseudo-F | P(perm) | U. perms | P(MC) | ECV (%) |
|---|---|---|---|---|---|---|---|---|
| Depth | 1 | 243.24 | 243.24 | 26.059 | 0.1045 | 10 | **0.0084** | 50.6 |
| Year | 2 | 4.0332 | 2.0166 | 0.46078 | 0.6431 | 9956 | 0.658 | 0.0 |
| Quadrat(Depth) | 4 | 37.336 | 9.3341 | 2.1328 | 0.1677 | 9965 | 0.1668 | 12.8 |
| DepthxYear | 2 | 24.211 | 12.106 | 2.766 | 0.1236 | 9957 | 0.1231 | 15.9 |
| Residual | 8 | 35.012 | 4.3765 | | | | | 20.8 |
| Total | 17 | 343.83 | | | | | | |

***E) Great Barrier Reef: Great Detached***

| Source | df | SS | MS | Pseudo-F | P(perm) | U. perms | P(MC) | ECV (%) |
|---|---|---|---|---|---|---|---|---|
| Depth | 1 | 24.818 | 24.818 | 2.5049 | 0.4044 | 10 | 0.1943 | 22.9 |
| Year | 2 | 9.0911 | 4.5456 | 3.889 | 0.0682 | 9967 | 0.0664 | 13.3 |
| Quadrat(Depth) | 4 | 39.632 | 9.9079 | 8.4768 | **0.0058** | 9948 | **0.0052** | 30.4 |
| DepthxYear | 2 | 6.1295 | 3.0647 | 2.6221 | 0.1387 | 9957 | 0.1346 | 14.1 |
| Residual | 8 | 9.3506 | 1.1688 | | | | | 19.2 |
| Total | 17 | 89.021 | | | | | | |

#### *F) Great Barrier Reef: Tijou*

| Source | df | SS | MS | Pseudo-F | P(perm) | U. perms | P(MC) | ECV (%) |
|---|---|---|---|---|---|---|---|---|
| Depth | 1 | 125.11 | 125.11 | 15.223 | 0.1001 | 10 | **0.0148** | 55.1 |
| Year | 2 | 2.2363 | 1.1182 | 2.7472 | 0.1303 | 9956 | 0.1272 | 5.3 |
| Quadrat(Depth) | 4 | 32.874 | 8.2186 | 20.193 | **0.0005** | 9961 | **0.0003** | 24.7 |
| DepthxYear | 2 | 1.5032 | 0.75158 | 1.8466 | 0.2181 | 9945 | 0.217 | 5.2 |
| Residual | 8 | 3.2561 | 0.40701 | | | | | 9.8 |
| Total | 17 | 164.98 | | | | | | |

#### *G) Great Barrier Reef: Yonge*

| Source | df | SS | MS | Pseudo-F | P(perm) | U. perms | P(MC) | ECV (%) |
|---|---|---|---|---|---|---|---|---|
| Depth | 1 | 129.62 | 129.62 | 13.113 | 0.0994 | 10 | **0.0366** | 54.7 |
| Year | 1 | 5.5799 | 5.5799 | 4.757 | 0.1185 | 9144 | 0.1175 | 10.5 |
| Quadrat(Depth) | 3 | 29.655 | 9.885 | 8.4272 | **0.0519** | 9357 | **0.0589** | 22.9 |
| DepthxYear | 1 | 0.30856 | 0.30856 | 0.26306 | 0.6423 | 9140 | 0.636 | 0.0 |
| Residual | 3 | 3.519 | 1.173 | | | | | 11.9 |
| Total | 9 | 168.38 | | | | | | |

**Table S5. Permutational analysis of variance (univariate PERMANOVA) on the change of scleractinian coral coverage (2012 vs. 2016/2017) by site and pairwise comparisons.** Test based on Euclidean distances on fourth root transformed and standardized data. Significant differences are indicated in bold. P(perm): *P*-value based on permutations, U. perms: Unique permutations, P(MC): Monte Carlo *P*- value, ECV(%): Estimated components of variation.

#### *A) Coral Sea: Osprey – Dutch Towers*

| Source | df | SS | MS | Pseudo-F | P(perm) | U. perms | P(MC) | ECV (%) |
|---|---|---|---|---|---|---|---|---|
| Depth | 1 | 41.507 | 41.507 | 11.445 | 0.1039 | 10 | **0.0266** | 40.6 |
| Year | 1 | 8.7938 | 8.7938 | 61.544 | **0.003** | 9472 | **0.0015** | 19.4 |
| Quadrat(Depth) | 4 | 14.507 | 3.6267 | 25.382 | **0.0065** | 9935 | **0.0045** | 21.3 |
| DepthxYear | 1 | 1.9507 | 1.9507 | 13.652 | **0.0262** | 9426 | **0.0219** | 12.5 |
| Residual | 4 | 0.57154 | 0.14289 | | | | | 6.1 |
| Total | 11 | 67.329 | | | | | | |

*DepthxYear*

| Depth | Groups | t | P(perm) | U. perms | P(MC) |
|---|---|---|---|---|---|
| 10 m | 2012, 2017 | 4.7886 | 0.1006 | 10 | **0.0099** |
| 40 m | 2012, 2017 | 2.2891 | 0.2076 | 10 | 0.0844 |

***B) Coral Sea: Osprey – Bigeye Ledge***

| Source | df | SS | MS | Pseudo-F | P(perm) | U. perms | P(MC) | ECV (%) |
|---|---|---|---|---|---|---|---|---|
| Depth | 1 | 19.61 | 19.61 | 1.1326 | 0.3967 | 10 | 0.3506 | 9.2 |
| Year | 1 | 8.0154 | 8.0154 | 9.5027 | **0.0398** | 9604 | **0.0377** | 16.3 |
| Quadrat(Depth) | 4 | 69.254 | 17.314 | 20.526 | **0.0091** | 9924 | **0.0057** | 42.9 |
| DepthxYear | 1 | 5.0908 | 5.0908 | 6.0355 | 0.0728 | 9571 | 0.0662 | 17.8 |
| Residual | 4 | 3.3739 | 0.84348 | | | | | 13.7 |
| Total | 11 | 105.34 | | | | | | |

***C) Coral Sea: Osprey – Nautilus Wall***

| Source | df | SS | MS | Pseudo-F | P(perm) | U. perms | P(MC) | ECV (%) |
|---|---|---|---|---|---|---|---|---|
| Depth | 1 | 19.61 | 19.61 | 1.1326 | 0.4127 | 10 | 0.3441 | 9.2 |
| Year | 1 | 8.0154 | 8.0154 | 9.5027 | **0.0395** | 9593 | **0.0378** | 16.3 |
| Quadrat(Depth) | 4 | 69.254 | 17.314 | 20.526 | **0.0082** | 9929 | **0.0061** | 42.9 |
| DepthxYear | 1 | 5.0908 | 5.0908 | 6.0355 | 0.0744 | 9571 | 0.0704 | 17.8 |
| Residual | 4 | 3.3739 | 0.84348 | | | | | 13.7 |
| Total | 11 | 105.34 | | | | | | |

***D) Coral Sea: Osprey – Holmes***

| Source | df | SS | MS | Pseudo-F | P(perm) | U. perms | P(MC) | ECV (%) |
|---|---|---|---|---|---|---|---|---|
| Depth | 1 | 160.34 | 160.34 | 20.138 | 0.1047 | 10 | **0.0122** | 45.8 |
| Year | 1 | 4.0286 | 4.0286 | 0.78073 | 0.4207 | 9511 | 0.4312 | 0.0 |
| Quadrat(Depth) | 4 | 31.849 | 7.9621 | 1.543 | 0.3448 | 9935 | 0.3372 | 10.7 |
| DepthxYear | 1 | 24.196 | 24.196 | 4.6891 | 0.1044 | 9630 | 0.0949 | 22.9 |
| Residual | 4 | 20.64 | 5.16 | | | | | 20.6 |
| Total | 11 | 241.05 | | | | | | |

***E) Great Barrier Reef: Great Detached***

| Source | df | SS | MS | Pseudo-F | P(perm) | U. perms | P(MC) | ECV (%) |
|---|---|---|---|---|---|---|---|---|
| Depth | 1 | 15.834 | 15.834 | 2.0114 | 0.4044 | 10 | 0.2276 | 18.0 |
| Year | 1 | 8.4553 | 8.4553 | 9.2571 | **0.046** | 9452 | **0.0358** | 17.5 |
| Quadrat(Depth) | 4 | 31.489 | 7.8723 | 8.6188 | **0.0353** | 9922 | **0.0309** | 29.1 |
| DepthxYear | 1 | 6.106 | 6.106 | 6.6851 | 0.0651 | 9612 | 0.0591 | 20.5 |
| Residual | 4 | 3.6535 | 0.91338 | | | | | 14.9 |
| Total | 11 | 65.538 | | | | | | |

***F) Great Barrier Reef: Tijou***

| Source | df | SS | MS | Pseudo-F | P(perm) | U. perms | P(MC) | ECV (%) |
|---|---|---|---|---|---|---|---|---|
| Depth | 1 | 89.646 | 89.646 | 17.479 | 0.1022 | 10 | **0.0142** | 53.9 |
| Year | 1 | 2.0679 | 2.0679 | 4.0431 | 0.1205 | 9574 | 0.1156 | 7.3 |
| Quadrat(Depth) | 4 | 20.515 | 5.1288 | 10.028 | **0.0299** | 9923 | **0.022** | 21.8 |
| DepthxYear | 1 | 1.1656 | 1.1656 | 2.2789 | 0.2064 | 9493 | 0.2003 | 6.7 |
| Residual | 4 | 2.0458 | 0.51146 | | | | | 10.3 |
| Total | 11 | 115.44 | | | | | | |

**Table S6. Permutational analysis of variance (univariate PERMANOVA) on the relative change in scleractinian coral coverage by site and pairwise comparisons between *Years* and the interaction *DepthxYear*.** Test based on Euclidean distances. Significant differences are indicated in bold. P(perm): *P*-value based on permutations, U. perms: Unique permutations, P(MC): Monte Carlo *P*- value, ECV(%): Estimated components of variation.

***A) Coral Sea: Osprey – Dutch Towers***

| Source | df | SS | MS | Pseudo-F | P(perm) | U. perms | P(MC) | ECV (%) |
|---|---|---|---|---|---|---|---|---|
| Depth | 1 | 631.34 | 631.34 | 2.2004 | 0.2983 | 10 | 0.2075 | 13.9 |
| Year | 2 | 2251.7 | 1125.9 | 15.011 | **0.0017** | 9942 | **0.0021** | 29.6 |
| Quadrat(Depth) | 4 | 1147.7 | 286.91 | 3.8254 | **0.0482** | 9964 | 0.0549 | 18.8 |
| DepthxYear | 2 | 549.07 | 274.54 | 3.6604 | 0.073 | 9948 | 0.069 | 18.3 |
| Residual | 8 | 600.02 | 75.003 | | | | | 19.4 |
| Total | 17 | 5179.8 | | | | | | |

*Year*

| Groups | t | P(perm) | U. perms | P(MC) |
|---|---|---|---|---|
| 2013, 2015 | 3.3944 | **0.0301** | 9611 | **0.0308** |
| 2013, 2017 | 4.3888 | **0.0136** | 9591 | **0.0128** |
| 2015, 2017 | 3.2174 | **0.0374** | 9529 | **0.0302** |

***B) Coral Sea: Osprey – Bigeye Ledge***

| Source | df | SS | MS | Pseudo-F | P(perm) | U. perms | P(MC) | ECV (%) |
|---|---|---|---|---|---|---|---|---|
| Depth | 1 | 769.42 | 769.42 | 3.3263 | 0.2011 | 10 | 0.141 | 14.8 |
| Year | 2 | 2183.9 | 1092 | 8.4162 | **0.0107** | 9957 | **0.0108** | 24.3 |
| Quadrat(Depth) | 4 | 925.25 | 231.31 | 1.7828 | 0.2279 | 9957 | 0.2255 | 11.1 |
| DepthxYear | 2 | 1541.3 | 770.63 | 5.9396 | **0.0286** | 9957 | **0.0246** | 28.0 |
| Residual | 8 | 1038 | 129.74 | | | | | 21.8 |
| Total | 17 | 6457.8 | | | | | | |

*DepthxYear by Depth*

| Depth | Groups | t | P(perm) | U. perms | P(MC) |
|---|---|---|---|---|---|
| 10 m | 2013, 2015 | 1.2308 | 0.2899 | 38 | 0.3425 |
| | 2013, 2017 | 3.8611 | 0.1024 | 38 | 0.0639 |
| | 2015, 2017 | 4.8601 | 0.1013 | 38 | **0.0432** |
| 40 m | 2013, 2015 | 1.278 | 0.3988 | 38 | 0.3278 |
| | 2013, 2017 | 0.74405 | 0.5996 | 16 | 0.5275 |
| | 2015, 2017 | 0.029295 | 1 | 38 | 0.9804 |

*DepthxYear by Year*

| Year | Groups | t | P(perm) | U. perms | P(MC) |
|---|---|---|---|---|---|
| 2013 | 10m, 40m | 0.36069 | 0.8021 | 10 | 0.7458 |
| 2015 | | 0.16819 | 1 | 10 | 0.8728 |
| 2017 | | 2.7588 | 0.099 | 10 | 0.0528 |

*Year*

| Groups | t | P(perm) | U. perms | P(MC) |
|---|---|---|---|---|
| 2013, 2015 | 1.7134 | 0.1621 | 9556 | 0.1657 |
| 2013, 2017 | 3.4646 | **0.0298** | 9595 | **0.0271** |
| 2015, 2017 | 2.5034 | 0.0656 | 9609 | 0.0671 |

#### *C) Coral Sea: Osprey – Nautilus Wall*

| Source | df | SS | MS | Pseudo-F | P(perm) | U. perms | P(MC) | ECV (%) |
|---|---|---|---|---|---|---|---|---|
| Depth | 1 | 2700.8 | 2700.8 | 32.717 | 0.1007 | 10 | **0.0054** | 28.1 |
| Year | 2 | 1721.6 | 860.82 | 6.1285 | **0.0247** | 9962 | **0.0239** | 18.1 |
| Quadrat(Depth) | 4 | 330.21 | 82.551 | 0.58772 | 0.6806 | 9966 | 0.6915 | 0.0 |
| DepthxYear | 2 | 2873.1 | 1436.6 | 10.227 | **0.0062** | 9950 | **0.006** | 34.3 |
| Residual | 8 | 1123.7 | 140.46 | | | | | 19.5 |
| Total | 17 | 8749.4 | | | | | | |

*DepthxYear by Depth*

| Depth | Groups | t | P(perm) | U. perms | P(MC) |
|---|---|---|---|---|---|
| 10 m | 2013, 2015 | 2.7757 | 0.166 | 48 | 0.1086 |
| | 2013, 2017 | 4.3736 | 0.0989 | 38 | **0.0452** |
| | 2015, 2017 | 3.5272 | 0.1003 | 38 | 0.0674 |
| 40 m | 2013, 2015 | 1.2663 | 0.3462 | 38 | 0.3383 |
| | 2013, 2017 | 0.70631 | 0.5336 | 38 | 0.5594 |
| | 2015, 2017 | 0.23875 | 1 | 38 | 0.8329 |

*DepthxYear by Year*

| Year | Groups | t | P(perm) | U. perms | P(MC) |
|---|---|---|---|---|---|
| 2013 | 10m, 40m | 0.12799 | 1 | 10 | 0.9003 |
| 2015 | | 2.1417 | 0.2041 | 10 | 0.0961 |
| 2017 | | 5.0695 | 0.1009 | 10 | **0.0072** |

*Year*

| Groups | t | P(perm) | U. perms | P(MC) |
|---|---|---|---|---|
| 2013, 2015 | 0.54802 | 0.616 | 9668 | 0.6079 |
| 2013, 2017 | 2.2776 | 0.0839 | 9536 | 0.0884 |
| 2015, 2017 | 2.9059 | **0.0486** | 9629 | **0.0433** |

#### *D) Coral Sea: Osprey – Holmes*

| Source | df | SS | MS | Pseudo-F | P(perm) | U. perms | P(MC) | ECV (%) |
|---|---|---|---|---|---|---|---|---|
| Depth | 1 | 980.51 | 980.51 | 1.218 | 0.3992 | 10 | 0.3329 | 12.1 |
| Year | 1 | 524.48 | 524.48 | 2.2649 | 0.2106 | 9624 | 0.2081 | 15.7 |
| Quadrat(Depth) | 4 | 3220.1 | 805.04 | 3.4764 | 0.1408 | 9930 | 0.133 | 38.0 |
| DepthxYear | 1 | 1.6211 | 1.6211 | 0.0070006 | 0.9294 | 9628 | 0.9346 | 0.0 |
| Residual | 4 | 926.28 | 231.57 | | | | | 34.2 |
| Total | 11 | 5653 | | | | | | |

#### *E) Great Barrier Reef: Great Detached*

| Source | df | SS | MS | Pseudo-F | P(perm) | U. perms | P(MC) | ECV (%) |
|---|---|---|---|---|---|---|---|---|
| Depth | 1 | 3647.2 | 3647.2 | 13.436 | 0.1022 | 10 | **0.0239** | 42.5 |
| Year | 1 | 131.54 | 131.54 | 1.9657 | 0.2337 | 9542 | 0.2342 | 5.9 |
| Quadrat(Depth) | 4 | 1085.8 | 271.44 | 4.0563 | 0.079 | 9936 | 0.1057 | 18.1 |
| DepthxYear | 1 | 396.34 | 396.34 | 5.9226 | 0.0687 | 9643 | 0.0746 | 18.8 |
| Residual | 4 | 267.68 | 66.92 | | | | | 14.7 |
| Total | 11 | 5528.5 | | | | | | |

#### *F) Great Barrier Reef: Tijou*

| Source | df | SS | MS | Pseudo-F | P(perm) | U. perms | P(MC) | ECV (%) |
|---|---|---|---|---|---|---|---|---|
| Depth | 1 | 3292.9 | 3292.9 | 1.8124 | 0.2978 | 10 | 0.2507 | 24.4 |
| Year | 1 | 550.47 | 550.47 | 4.6205 | 0.1004 | 9621 | 0.0968 | 13.2 |
| Quadrat(Depth) | 4 | 7267.3 | 1816.8 | 15.25 | **0.0118** | 9937 | **0.0095** | 45.4 |
| DepthxYear | 1 | 11.276 | 11.276 | 0.094646 | 0.7113 | 9324 | 0.777 | 0.0 |
| Residual | 4 | 476.54 | 119.14 | | | | | 17.0 |
| Total | 11 | 11598 | | | | | | |

**Table S7. Permutational analysis of variance (PERMANOVA) on the structure of scleractinian coral communities by family.** Test based on Bray-Curtis distances on fourth root transformed and standardized data. Significant differences are indicated in bold. P(perm): *P*-value based on permutations, U. perms: Unique permutations, P(MC): Monte Carlo *P*- value, ECV(%): Estimated components of variation.

| Source | df | SS | MS | Pseudo-F | P(perm) | U. perms | P(MC) | ECV (%) |
|---|---|---|---|---|---|---|---|---|
| Region | 1 | 592.74 | 592.74 | 0.76499 | 0.5563 | 9964 | 0.5338 | 0.0 |
| Depth | 1 | 5454.4 | 5454.4 | 5.6834 | **0.0054** | 9966 | **0.0107** | 14.7 |
| Year | 5 | 785.76 | 157.15 | 1.4218 | 0.2387 | 9947 | 0.2557 | 2.4 |
| Site(Region) | 5 | 11865 | 2373 | 3.2215 | **0.0002** | 9919 | **0.0006** | 14.2 |
| RegionxDepth | 1 | 793.83 | 793.83 | 1.8405 | 0.1883 | 9967 | 0.186 | 9.3 |
| RegionxYear** | 0 | 0 | | No test | | | | 0.0 |
| DepthxYear | 5 | 1001.4 | 200.28 | 2.1466 | 0.0734 | 9942 | 0.0745 | 5.1 |
| Site(Region)xDepth | 5 | 6209.6 | 1241.9 | 1.686 | 0.0573 | 9923 | 0.0734 | 11.2 |
| Site(Region)xYear** | 11 | 1216.9 | 110.63 | 1.5966 | 0.0644 | 9918 | 0.0675 | 4.0 |
| RegionxDepthxYear** | 0 | 0 | | No test | | | | 0.0 |
| Quadrat(Site(Region)xDepth) | 27 | 20639 | 764.41 | 11.032 | **0.0001** | 9858 | **0.0001** | 22.0 |
| Site(Region)xDepthxYear** | 11 | 1027 | 93.36 | 1.3474 | 0.1718 | 9930 | 0.1649 | 4.3 |
| Residual | 63 | 4365.1 | 69.288 | | | | | 12.7 |
| Total | 135 | 61661 | | | | | | |

**Table S8. Permutational analysis of variance (PERMANOVA) on the structure of scleractinian coral communities by morphologies.** Test based on Bray-Curtis distances on fourth root transformed and standardized data. Significant differences are indicated in bold. P(perm): *P*-value based on permutations, U. perms: Unique permutations, P(MC): Monte Carlo *P*- value, ECV(%): Estimated components of variation.

| Source | df | SS | MS | Pseudo-F | P(perm) | U. perms | P(MC) | ECV (%) |
|---|---|---|---|---|---|---|---|---|
| Region | 1 | 930.7 | 930.7 | 1.5819 | 0.2325 | 9964 | 0.2518 | 6.5 |
| Depth | 1 | 6737.3 | 6737.3 | 5.0131 | **0.0183** | 9967 | **0.0213** | 16.5 |
| Year | 5 | 1908.3 | 381.65 | 10.323 | **0.0002** | 9935 | **0.001** | 6.7 |
| Site(Region) | 5 | 8817.7 | 1763.5 | 3.1671 | **0.0012** | 9935 | **0.0012** | 12.5 |
| RegionxDepth | 1 | 436.72 | 436.72 | 0.7431 | 0.5624 | 9961 | 0.5626 | 0.0 |
| RegionxYear** | 0 | 0 | | No test | | | | 0.0 |
| DepthxYear | 5 | 624.35 | 124.87 | 1.1276 | 0.3894 | 9943 | 0.4105 | 1.9 |
| Site(Region)xDepth | 5 | 8806.9 | 1761.4 | 3.1633 | **0.0003** | 9909 | **0.0009** | 17.7 |
| Site(Region)xYear** | 11 | 405.8 | 36.891 | 0.53011 | 0.8901 | 9933 | 0.8858 | 0.0 |
| RegionxDepthxYear** | 0 | 0 | | No test | | | | 0.0 |
| Quadrat(Site(Region)xDepth) | 27 | 15582 | 577.13 | 8.2931 | **0.0001** | 9856 | **0.0001** | 19.3 |
| Site(Region)xDepthxYear** | 11 | 1219.2 | 110.84 | 1.5927 | 0.079 | 9925 | 0.0832 | 5.8 |
| Residual | 63 | 4384.2 | 69.591 | | | | | 13.0 |
| Total | 135 | 58369 | | | | | | |

**Table S9. Permutational analysis of variance (PERMANOVA) on the structure of benthic communities.** Test based on Bray-Curtis distances on fourth root transformed and standardized data. Significant differences are indicated in bold. P(perm): *P*-value based on permutations, U. perms: Unique permutations, P(MC): Monte Carlo *P*- value, ECV(%): Estimated components of variation.

| Source | df | SS | MS | Pseudo-F | P(perm) | U. perms | P(MC) | ECV (%) |
|---|---|---|---|---|---|---|---|---|
| Region | 1 | 1134.7 | 1134.7 | 1.3991 | 0.2873 | 9959 | 0.2995 | 6.3 |
| Depth | 1 | 6475.5 | 6475.5 | 4.044 | **0.029** | 9943 | **0.0308** | 15.5 |
| Year | 5 | 1273.4 | 254.68 | 2.223 | **0.0236** | 9948 | 0.0575 | 4.2 |
| Site(Region) | 5 | 12896 | 2579.1 | 8.2768 | **0.0001** | 9910 | **0.0001** | 16.9 |
| RegionxDepth | 1 | 294.51 | 294.51 | 0.43788 | 0.7609 | 9963 | 0.7207 | 0.0 |
| RegionxYear** | 0 | 0 | | No test | | | | 0.0 |
| DepthxYear | 5 | 394.71 | 78.941 | 1.3544 | 0.2698 | 9946 | 0.3041 | 2.3 |
| Site(Region)xDepth | 5 | 10616 | 2123.2 | 6.8137 | **0.0001** | 9908 | **0.0001** | 21.4 |
| Site(Region)xYear** | 11 | 1262.3 | 114.76 | 3.1585 | **0.0002** | 9931 | **0.0003** | 5.6 |
| RegionxDepthxYear** | 0 | 0 | | No test | | | | 0.0 |
| Quadrat(Site(Region)xDepth) | 27 | 8723.2 | 323.08 | 8.8923 | **0.0001** | 9835 | **0.0001** | 14.3 |
| Site(Region)xDepthxYear** | 11 | 641.7 | 58.336 | 1.6056 | 0.0853 | 9931 | 0.0875 | 4.2 |
| Residual | 63 | 2289 | 36.333 | | | | | 9.3 |
| Total | 135 | 51910 | | | | | | |
